# Integrated hepatic ferroptosis gene signature dictates pathogenic features of ferroptosis

**DOI:** 10.1101/2024.10.27.620461

**Authors:** Takashi Matsumoto, Akihiro Nita, Yohei Kanamori, Ayato Maeda, Noriko Yasuda-Yoshihara, Kosuke Mima, Hirohisa Okabe, Katsunori Imai, Hiromitsu Hayashi, Yuta Matsuoka, Katsuya Nagaoka, Keiichi I. Nakayama, Yuki Sugiura, Yasuhito Tanaka, Hideo Baba, Toshiro Moroishi

## Abstract

**Background & Aims:** Ferroptosis, a distinctive form of cell death induced by iron-dependent lipid peroxidation, is implicated in various biological processes, including liver diseases. Establishing an iron overload-induced ferroptosis model and identifying hepatic gene signatures associated with ferroptosis are crucial for understanding its role in liver pathogenesis.

**Methods:** F-box and leucine-rich repeat protein 5 (FBXL5) is a substrate-recognition component of the SCF E3 ligase complex that restricts intracellular iron levels. In this study, we used liver-specific *Fbxl5*-null mice to establish an iron overload-induced ferroptosis model. Transcriptome analysis identified genes involved in hepatic ferroptosis. Integrating these gene signatures with another ferroptosis model enabled the assessment of ferroptosis-related pathology in murine liver injury models and in 174 patients undergoing liver resection surgery.

**Results:** Iron overload induced severe liver damage in liver-specific *Fbxl5*-null mice, characterized by elevated liver enzymes, histopathological changes, and lipid peroxidation. Transcriptome analysis revealed a distinct set of genes associated with hepatic ferroptosis response. Generating a gene signature for evaluating ferroptosis enhanced the understanding of ferroptosis-related pathologies in liver diseases. Iron overload exacerbated liver damage in murine ischemia-reperfusion injury models via ferroptosis induction. In human patients, elevated serum iron levels correlated with sustained post-operative liver damage, indicating heightened susceptibility to ferroptosis.

**Conclusion:** Here, a murine model of iron overload-induced hepatic ferroptosis was established, and a gene signature indicative of hepatic ferroptosis response in both mice and humans was identified. These findings underscore the role of ferroptosis in liver injury progression and suggest potential therapeutic targets for liver disease intervention.

**HIGHLIGHTS:**

- Liver-specific *Fbxl5* knockout mice provide an iron-induced ferroptotic injury model
- Integrated gene signature of iron- and acetaminophen-induced liver injury dictates ferroptosis
- Iron overload aggravates hepatic ischemia-reperfusion injury in mice
- Patients with high iron levels show delayed post-operative liver damage recovery

**IMPACT AND IMPLICATIONS:** Our study elucidated the critical role of iron in liver disease pathogenesis and ischemia-reperfusion injury (IRI). By establishing a murine model of iron overload-induced ferroptosis, we confirmed that iron overload exacerbated hepatic IRI, underscoring the importance of ferroptosis in liver damage. Additionally, the development of an integrated gene signature for hepatic ferroptosis response provides a valuable tool for evaluating ferroptosis in liver diseases. Via analysis of patient data, we also highlighted the clinical relevance of ferroptosis in post-operative liver damage, offering insights into potential therapeutic strategies targeting iron and ferroptosis to improve outcomes in patients with liver diseases.

## INTRODUCTION

Ferroptosis is a distinct form of cell death that differs from previously described modes, such as apoptosis, necrosis, and necroptosis, as it is induced by iron-dependent lipid peroxidation [1, 2]. Its morphological characteristics primarily include marked shrinkage of mitochondria, increased mitochondrial membrane density, and the reduction or disappearance of mitochondrial cristae [3]. Since its discovery, ferroptosis has been implicated in various biological contexts [4], including aging [5, 6], neurodegenerative disorders [7-9], infectious diseases [10, 11], autoimmune diseases [12, 13], and cancer [14-16]. Consistent with the broad biological impact of ferroptosis, emerging evidence indicates an association between ferroptosis and the pathogenesis of liver diseases, including liver injury, steatohepatitis, fibrosis, cirrhosis, and liver cancer [17]. The widespread availability of next-generation sequencing data in the public domain has fueled interest in developing gene signatures to monitor hepatic ferroptosis, thereby elucidating its connection to liver disease pathogenesis and evaluating its potential as a target for prevention and treatment. However, this has been challenging, particularly when identifying gene signatures that dictate tissue-level responses, because of the limited options for animal models for inducing hepatic ferroptosis.

Excess iron within cells can induce the Fenton reaction, producing highly reactive hydroxyl radicals that oxidizes phospholipids containing polyunsaturated fatty acids (PUFAs), leading to ferroptosis. Maintaining strict control over iron homeostasis and redox potential against oxidative stress is crucial for preventing ferroptosis, as imbalances can induce this form of cell death [18]. Ferroptosis can be initiated by a reduction in redox potential, such as the depletion of reduced glutathione (GSH) and decreased activity of glutathione peroxidase 4 (GPX4). Consequently, GSH depletion induced by an acetaminophen (APAP) overdose [19] and the conditional deletion of GPX4 in the liver [20] are widely used methods to induce hepatic ferroptosis. Despite the critical role of iron in ferroptosis induction, an iron overload-induced hepatic ferroptosis model has not yet been established.

In mammals, iron exists in two oxidation states, ferrous iron (Fe^2+^) and ferric iron (Fe^3+^). Iron absorbed by cells are released into the cytosol in its ferrous form. Ferrous iron is either directly integrated into iron-dependent proteins or directed to the mitochondria, where it becomes part of heme and iron–sulfur clusters [21]. However, ferrous iron is redox-active and triggers the generation of reactive oxygen species. To mitigate its harmful effects, any unused or excess ferrous iron is converted into the ferric form and stored within ferritin [22, 23]. Ferric iron is relatively safer in terms of oxidative damage but must be reduced back to the ferrous state to enter the cellular bioavailable iron pool. Because of the distinct biological characteristics of ferrous and ferric iron, their valence states and quantities influence both biological and pathological outcomes related to iron. Ferrous iron is considered a potent inducer of ferroptosis. Cellular levels of ferrous iron are primarily regulated *in vivo* by F-box and leucine-rich repeat protein 5 (FBXL5) [24]. FBXL5 is a substrate-recognition component of the SCF E3 ligase complex that mediates the ubiquitination and degradation of iron regulatory protein 1 (IRP1, also known as ACO1) and IRP2 (also known as IREB2) [25, 26], thus controlling intracellular ferrous iron levels by regulating the protein expression of iron metabolism-related components at the post-transcriptional level. We have previously reported that the loss of FBXL5 in mice results in embryonic lethality, whereas liver-specific *Fbxl5* ablation is associated with ferrous iron accumulation and mitochondrial dysfunction in hepatocytes, leading to steatohepatitis [24]. Additionally, when these mice were fed a high-iron diet, they died of acute liver failure due to excess iron. In the present study, we used this animal model to establish an iron overload-induced ferroptosis model and identify commonly regulated genes in the hepatic response to ferroptosis using a transcriptome-wide approach. We used this newly generated gene set to uncover the key functions of iron in the pathogenesis of hepatic ischemia-reperfusion injury (IRI) in both mice and humans.

## MATERIALS AND METHODS

### Mice

*Fbxl5*^F/F^ mice were generated as described previously [24]. These mice were crossed with *Alb*-Cre transgenic mice [27] to generate *Alb*-Cre/*Fbxl5*^F/F^ mice. Age- (8–14 weeks) and sex-matched littermates were used for the experiments. All mice were backcrossed with C57BL/6 mice for more than seven generations under specific pathogen-free conditions. All mouse experiments were approved by the Animal Ethics Committee of Kumamoto University (A2021-061).

### Patients data

Tissue specimens were obtained from patients with hepatocellular carcinoma who underwent hepatectomy at the Department of Gastroenterological Surgery, Kumamoto University Hospital, between April 2007 and October 2023 (n = 1,014 patients). Patients were excluded if pre-operative serum iron levels were not measured (n = 840). Finally, 174 patients were retrospectively analyzed. For the data on ischemic time, eight patients were excluded because the exact duration of hepatic ischemia performed using the Pringle maneuver method was not clear from the operation records, leaving 166 patients for final analysis. The present study was approved by the Medical Ethics Committee of Kumamoto University (Project No. 1291 and 2107), and written informed consent was obtained from all participants. The specimens of the cancerous tissue and adjacent background tissue of the surgical resection specimen were collected as a single mass, embedded in an optimal cutting temperature (O.C.T.) compound, quickly frozen on dry ice, and stored at −80L. The frozen specimens were sliced into 20-micrometer thin sections, the background non-cancer areas were identified by H&E staining, and RNA of this area was separately extracted using the RNeasy mini kit (QIAGEN, Cat# 74104).

### Iron administration

The animals were restrained by tight scruffing using a non-gavage hand. Oral gavage was performed using 1.5 inch, 20-gauge, stainless steel feeding needles with a 1.9 mm ball (Sigma-Aldrich, Cat# CAD9921), administering a final dose of 600 mg/kg of ferric ammonium citrate (FAC; Sigma-Aldrich, Cat# M0512) in 15 mL/kg of gavage solution. The gavage solution consisted of a mixture of 67% olive oil (Fujifilm, Cat# 150-00276) and 33% water containing 120 mg/mL of FAC (resulting in a final concentration of 40 mg/mL FAC in the gavage solution). A vehicle solution comprising 67% olive oil and 33% water was used as the control. For mass spectrometry experiments, the mice were administered a final dose of 1,500 mg/kg FAC via oral gavage. Iron gavage was administered after 12–16 h of fasting. For damage prevention experiments, the ferroptosis inhibitor ferrostatin-1 (Fer-1; Cayman Chemical, Cat# 17729) was dissolved in a vehicle buffer containing 10% dimethyl sulfoxide (DMSO; Fujifilm, Cat# 046-21981), 45% polyethylene glycol 300 (PEG300; Selleck, Cat# S6704), and 45% phosphate-buffered saline (PBS). Fer-1 solution or vehicle was administered intraperitoneally 1 h before the oral gavage experiment.

### Liquid chromatography-high resolution tandem mass spectrometry (LC-HRMS/MS) analysis of oxidized phosphatidylcholines (PC)

LC-MS/MS analysis was performed using a Nexera LC system (Shimadzu Co., Kyoto, Japan) coupled with a high-performance benchtop quadrupole Orbitrap mass spectrometer (Q Exactive; Thermo Fisher Scientific, Waltham, MA, USA) equipped with an electrospray ionization source. The LC parameters, ionization conditions, and experimental conditions for MS/MS and parallel reaction monitoring have been described previously [28]. LC-HRMS/MS analysis was performed using Xcalibur 4.2.47 software (Thermo Fisher Scientific).

### Matrix-assisted laser desorption/ionization-tandem MS-MS imaging (MALDI-MS/MS/MSI) of oxidized lipids

Sample preparation for MALDI-MS/MS/MSI was performed as described previously [28]. Briefly, thin liver sections (8 µm) were prepared using a cryomicrotome (CM3050; Leica Microsystems, Tokyo, Japan) and mounted on indium tin oxide-coated glass slides (Bruker Daltonics GmbH, Leipzig, Germany). The slides were coated with 2,5-dihydroxybenzoic acid solution (50Lmg/mL in 80% aqueous ethanol) by manual spraying using an artistic brush (Procon Boy FWA Platinum; Mr. Hobby, Tokyo, Japan). MALDI-MS/MS and MALDI-MSI experiments were performed using a MALDI linear ion trap mass spectrometer (MALDI LTQ XL; Thermo Fisher Scientific, Bremen, Germany) equipped with a 60-Hz N2 laser (λL=L337Lnm). The laser energy and the raster step size were set at 32 μJ and 60 µm, respectively. Auto-gain control (in which the ion trap is filled with the optimum number of ions) was not utilized in this experiment. Mass spectra were acquired in positive ion mode, and MS/MS transitions for PC PUFA;^18^O2 imaging were as follows: m/z 794.5L→L774.5 (neutral loss of H_2_^18^O) for PC34:2;^18^O2 with a precursor ion isolation width of m/z 1.0. After sample analysis, ion images were reconstructed using the ImageQuest v. 1.0.1 software (Thermo Fisher Scientific).

### Mouse models for IRI

Mice were fasted for 12–16 h prior to the operation, followed by anesthesia with medetomidine (10 mg/kg), midazolam (50 mg/kg), and butorphanol (50 mg/kg) via intraperitoneal injection. A midline incision was made, and an atraumatic clip was placed across the portal vein, hepatic artery, and bile duct just above the right branch. This procedure interrupts blood flow to the left lateral and median lobes, representing approximately 70% of the total blood supply to the liver. After 30 min of partial hepatic ischemia, the clip was removed to initiate reperfusion. Sham mice underwent the same procedure without vascular occlusion. After 3 h of reperfusion, the mice were euthanized via exsanguination under anesthesia and serum samples were collected. The left lateral and median lobes of the liver were collected and immediately fixed in 4% phorbol myristate acetate and stored at **−**80 L until further analysis. For damage prevention experiments, Fer-1 was dissolved in vehicle buffer containing 10% DMSO, 45% PEG300, and 45% PBS. Fer-1 solution or vehicle was administered intraperitoneally 1 h before IRI.

### Reverse transcription (RT) and real-time polymerase chain reaction (PCR) analysis

Total RNA was isolated from the liver using ISOGEN (NIPPON GENE, Cat#319-90211) and reverse-transcribed to complementary DNA (cDNA) using PrimeScript RT Master Mix (Takara, Cat# RR036A). The cDNA was then diluted and used for quantification by performing real-time PCR in a StepOnePlus Real-Time PCR System (Thermo Fisher Scientific) with the SYBR Premix Ex Taq II kit (Takara, Cat# RR820). The PCR primers used were as follows:

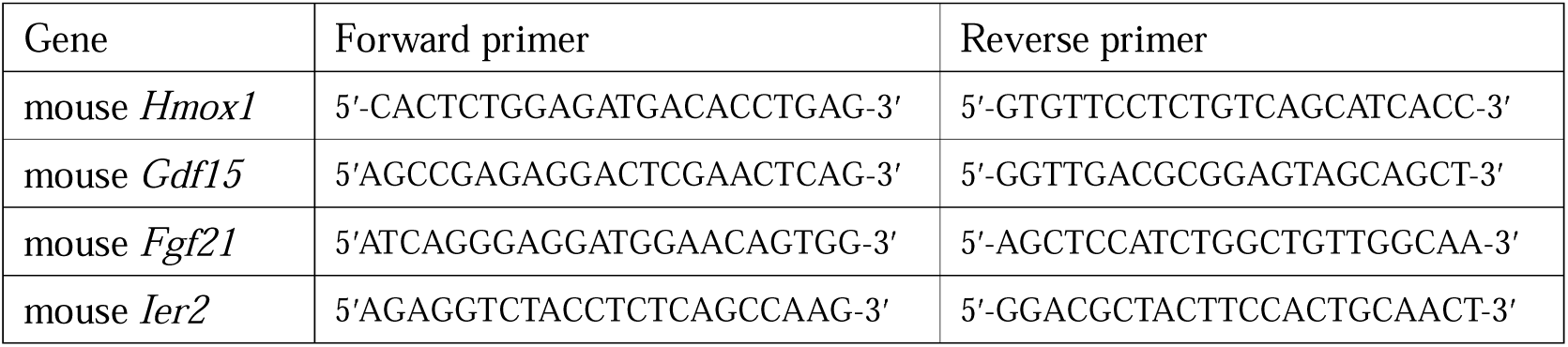

### RNA sequencing and data analysis

Total RNA was extracted from the livers of fasting mice using TRIzol (Life Technologies, Cat#15596-018) according to the manufacturer’s instructions. The RNA was then subjected to sequencing using an Illumina NovaSeq6000 sequencer (2 × 150 bp read length, San Diego, CA, USA). The quality of the raw paired-end reads was assessed using FastQC (version v0.12.1), and adaptor sequences were trimmed using Trim Galore (version 0.6.10). Paired-end reads were mapped against mouse (GRCm39) or human (GRCh38) genomes and analyzed using a series of programs, including HISAT2 (version 2.1.1), featureCounts (version 2.16.1), and DESeq2 (version 1.42.1). The expression data were further analyzed using GSEA v4.1.0 software. Gene sets for gene set enrichment analysis (GSEA) were obtained from the Molecular Signatures Database v4.0, which is available on the GSEA website. Raw RNA-seq data, supporting the findings of the present study, are available from the DNA Data Bank of Japan (DDBJ; accession numbers PRJDB18097 [mouse data] and PRJDB18207 [human data]).

### GSEA

The expression data were analyzed using GSEA v4.1.0 software. Gene sets for GSEA were obtained from the Molecular Signatures Database v4.0. The RNA sequencing data used for each figure are as follows: Fig. 3D, RNA sequencing data was obtained from GSE117066 [29], comparing normal group (n=3, indicated as control) and ischemic reperfusion group (n=3, subjected to 70% hepatic ischemic reperfusion); Fig. 3E, RNA sequencing data was obtained from Li et al. [30], comparing methionine–choline-deficient -diet group (n=3) and normal diet group (n=3); Fig. 3F: RNA sequencing data was obtained from GSE207855 [31], comparing control group (n=6) and carbon tetrachloride (CCl_4_) injection group (n=6); Fig. 4E: RNA sequencing data was obtained from GSE113024 [32], comparing donor liver at the end of preservation in conventional liver transplantation group (n=3) and post-reperfusion group (n=3).

### Histopathology and immunostaining

Histological analysis was performed as previously described [24], with slight modifications. Briefly, liver samples were fixed with 4% paraformaldehyde in PBS, embedded in paraffin, sectioned using a microtome, and stained with hematoxylin and eosin (H&E) according to standard protocols. For Perls’ Prussian blue staining to detect iron, deparaffinized and hydrated liver sections were incubated for 30 min at room temperature with Perls reagent (5% potassium ferrocyanide and 5% hydrogen chloride in deionized water) and then counterstained with hematoxylin solution for 30 s. For immunostaining analysis, deparaffinized and hydrated liver sections underwent antigen retrieval using HISTOFINE antigen retrieval solution (Nichirei, Cat# 415201). Tissues were then stained with antibodies against 4-hydroxynonenal (4-HNE; Abcam, Cat# ab46545) according to standard procedures and counterstained with hematoxylin solution for 30 s. Sections from control and mutant mice were processed simultaneously to monitor and detect nonspecific staining.

### Statistical analysis

All data were analyzed using GraphPad Prism (GraphPad Software, San Diego, CA, USA). mRNA analysis was performed using a base 10 logarithm transformation. Statistical parameters and methods are detailed in the figure legends. Statistical significance was defined as *p-*value < 0.05.

## RESULTS

### Establishment of an iron overload-induced ferroptosis model using liver-specific *Fbxl5*-null mice

We generated mice with liver-specific ablation of *Fbxl5* by crossing *Fbxl5*^F/F^ mice, carrying floxed *Fbxl5* alleles, with *albumin (Alb)-*Cre transgenic mice (expressing Cre recombinase under the regulation of the *Alb* promoter). To establish an iron overload-induced liver ferroptosis model, we conducted oral iron gavage experiments using liver-specific FBXL5-deficient [*Alb-Cre*/*Fbxl5*^F/F^; referred to as FBXL5 liver-KO (knockout)] mice and *Fbxl5*^F/F^ control mice (Fig. 1A). Although iron administration had minimal impact on control mice, the serum levels of aspartate aminotransferase (AST), alanine aminotransferase (ALT), and lactate dehydrogenase (LDH) dramatically increased in FBXL5 liver-KO mice at 3 and 6 h post-administration (Fig. 1B). Consistent with these results, microscopic analysis of H&E stained sections demonstrated increased hepatocyte damage in the livers of FBXL5 liver-KO mice (Fig. 1C). Perls iron staining further demonstrated increased iron accumulation in the hepatocytes of FBXL5 liver-KO mice compared to control mice (Fig. 1D). Immunostaining of 4-HNE-modified proteins indicated a marked increase in lipid peroxidation (Fig. 1E). Indeed, LC-HRMS/MS analysis of oxidized PC revealed a marked increase in single, double, and triple oxygenated PC PUFA levels (PC PUFA;O/O2/O3), characteristic of ferroptosis induction [33], in the livers of FBXL5 liver-KO mice compared to that in control mice upon iron administration (Fig. 1F). To investigate the association between oxidized PC PUFAs and hepatic cell death, we conducted *in vivo* ^18^O stable isotope labeling and MALDI-MS/MS/MSI to visualize endogenous oxidized PC PUFAs [28]. FBXL5 liver-KO mice were exposed to a flow of ^18^O_2_ air for 3Lh after iron administration. Subsequent analysis revealed significant accumulation of doubly oxygenated PC PUFAs, specifically PC34:2;^18^O2, in the damaged liver areas of FBXL5 liver-KO mice (Fig. 1G), suggesting an association between oxidized PC PUFA accumulation and liver damage induced by iron administration in these mice. To confirm that iron-induced hepatotoxicity in FBXL5 liver-KO mice was due to ferroptosis, we intraperitoneally injected Fer-1, a ferroptosis inhibitor, before iron administration. Treatment with Fer-1 reduced the levels of liver transaminases and LDH (Fig. 1H), histopathological liver damage (Fig. 1I), and levels of 4-HNE-modified proteins (Fig. 1J) compared to control DMSO treatment. Therefore, we established an *in vivo* mouse model of iron overload-induced ferroptosis by administering iron to FBXL5 liver-KO mice.

**Fig. 1.**
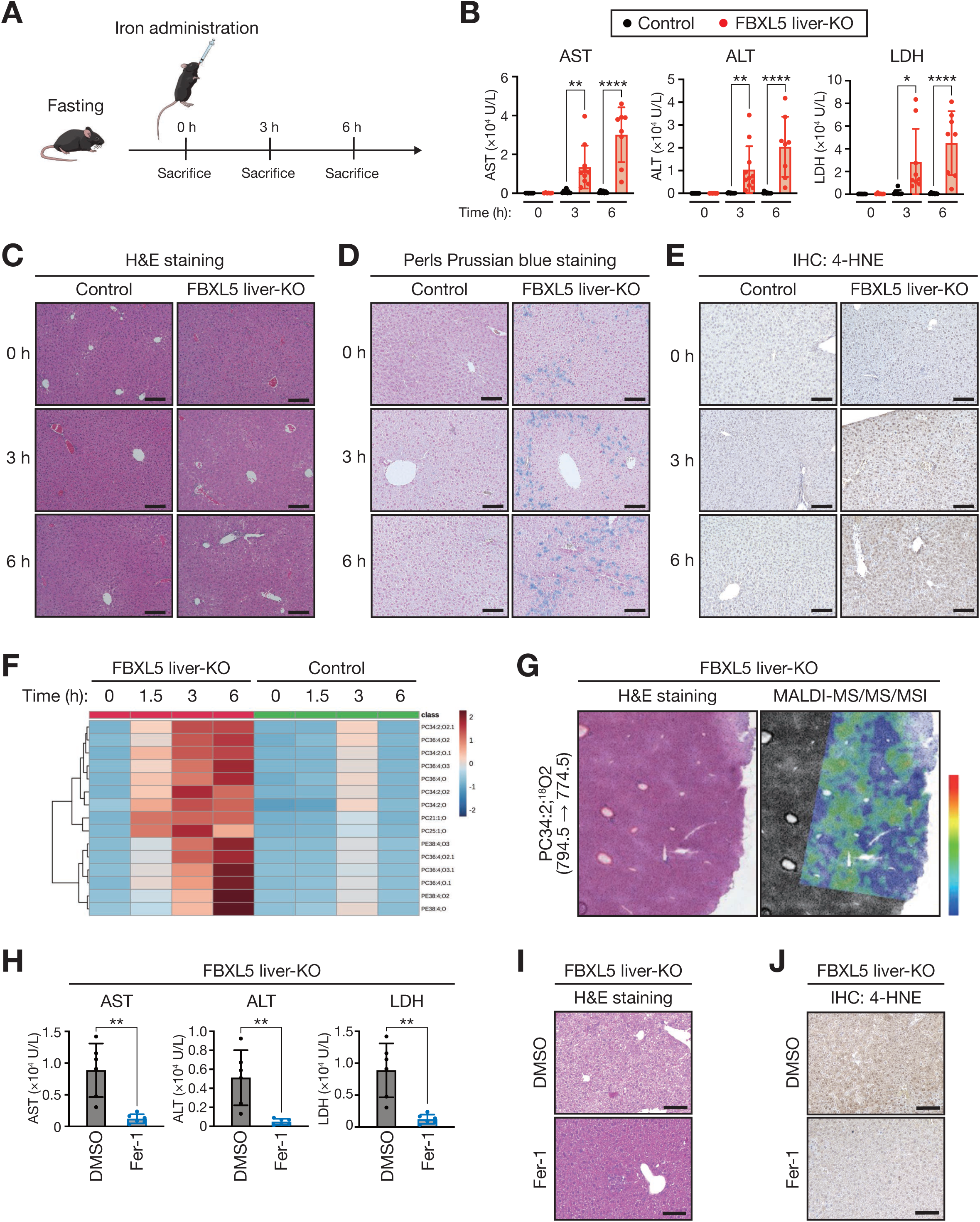
Establishment of an iron overload-induced ferroptosis model using liver-specific *Fbxl5*-null mice. (A) Schematic overview of the experimental procedure for ferroptosis induction. (B) Serum levels of aspartate transaminase (AST), alanine transaminase (ALT), and lactate dehydrogenase (LDH) in *Fbxl5*^F/F^ (control) and *Alb-*Cre*/Fbxl5*^F/F^ (FBXL5 liver-KO) mice at the indicated times after iron administration. Data are represented as mean ± standard deviation (SD). The number of mice used for this data are as follows: AST, n = 12, 8, and 7 mice at 0, 3, and 6 h, respectively, for the control group; and n = 6, 10, and 8 mice at 0, 3, and 6 h, respectively, for the FBXL5 liver-KO group; ALT, n = 12, 11, and 10 mice at 0, 3, and 6 h, respectively, for the control group; and n = 6, 11, 8 mice at 0, 3, 6 h, respectively, for the FBXL5 liver-KO group; LDH, n = 12, 9, and 7 mice at 0, 3, and 6 h, respectively, for the control group; and n = 6, 9, and 8 mice at 0, 3, and 6 h, respectively, for the FBXL5 liver-KO group. \**p* < 0.05; ∗∗*p* < 0.01; ∗∗∗∗*p* < 0.0001 [one-way analysis of variance (ANOVA) test followed by Tukey’s multiple comparison test]. (C-E) Histological analysis of the liver of *Alb-*Cre*/Fbxl5*^F/F^ (FBXL5 liver-KO) mice compared with that of *Fbxl5*^F/F^ (control) mice. Hematoxylin and eosin (H&E) staining is shown in (C), Perls Prussian blue staining for the detection of iron is shown in (D), immunohistochemical (IHC) staining with antibodies against 4-hydroxynonenal (4-HNE) is shown in (E). Scale bars represent 100 μm. (F) Heat map showing time-dependent profiles of endogenous oxidized phosphatidylcholines in lipid extracts from the liver of *Fbxl5*^F/F^ (control) and *Alb-*Cre*/Fbxl5*^F/F^ (FBXL5 liver-KO) mice at the indicated times after iron administration. The color reflects normalized mass spectrometry (MS) peak area of each oxidized phosphatidylcholines using log_10_ transformation and autoscaling. (G) Histological analysis of the liver of *Alb-*Cre*/Fbxl5*^F/F^ (FBXL5 liver-KO) mice was conducted at 3 h after iron administration. The H&E staining results are shown in the left panel, and the matrix-assisted laser desorption/ionization-tandem MS-MS imaging (MALDI-MS/MS/MSI) results for PC34:2;^18^O2 (794.5L→L774.5) are demonstrated in the right panel. (H) Serum levels of AST, ALT and LDH in *Alb-*Cre*/Fbxl5*^F/F^ (FBXL5 liver-KO) mice provided (or not) with the ferroptosis inhibitor ferrostatin-1 (Fer-1) for 1 h prior to iron exposure for 3 h. Data are represented as mean ± SD; n = 6 mice for each group. ∗∗*p* < 0.01 (unpaired t-test). (I and J) Histological analysis of the liver of *Alb-*Cre*/Fbxl5*^F/F^ (FBXL5 liver-KO) mice treated as in (H). H&E staining is shown in (I), IHC staining with antibodies against 4-HNE is shown in (J). Scale bars represent 100 μm.

### Liver-specific FBXL5 deficiency induces unique gene expression profiles upon iron administration

To investigate the changes in the liver transcriptome induced by ferroptosis, we examined the time-dependent gene expression profiles in the livers of FBXL5 liver-KO mice and control mice following iron administration (Fig. 2A and Table S1). Principal component analysis (Fig. 2B) and a heatmap of differentially expressed genes (DEGs) (Fig. 2C) revealed extensive alterations in the liver transcriptome of FBXL5 liver-KO mice at 3 and 6 h post iron administration. To investigate the immediate effects of ferroptosis on liver tissue response, we compared the gene expression patterns between FBXL5 liver-KO mice and control mice 3 h after iron administration (Table S2). GSEA highlighted the enrichment of ferroptosis-related gene set (mmu04216) in the livers of FBXL5 liver-KO mice (Fig. 2D), indicating the induction of ferroptosis. These findings collectively demonstrate that our mouse model of iron overload-induced ferroptosis can effectively replicate the pathogenesis of ferroptosis *in vivo*, characterized by both histological changes and alterations in gene expression profiles.

**Fig. 2.**
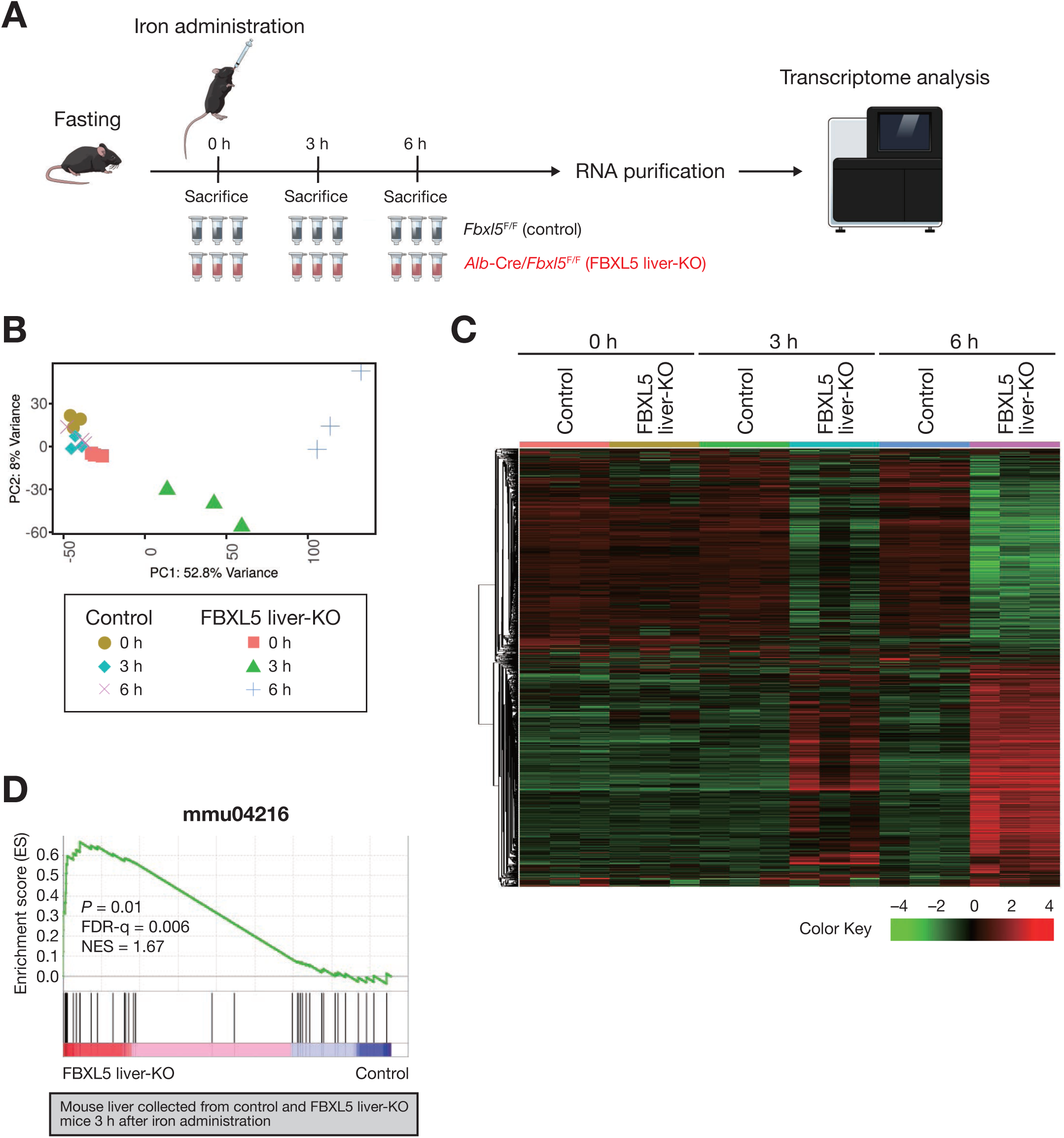
Characterization of the gene expression profiles of the liver in response to iron overload-induced ferroptosis. (A) Schematic overview of sample preparation for the transcriptome analysis. (B) Principal component analysis of the transcriptome of the liver of *Fbxl5*^F/F^ (control) and *Alb-*Cre*/Fbxl5*^F/F^ (FBXL5 liver-KO) mice at the indicated times after iron administration. (C) Heatmap correlation of *Fbxl5*^F/F^ (control) and *Alb-*Cre*/Fbxl5*^F/F^ (FBXL5 liver-KO) mice at the indicated times after iron administration. (D) Gene set enrichment analysis (GSEA) of the Kyoto Encyclopedia of Genes and Genomes (KEGG) ferroptosis pathway (mmu04216) in the liver of *Fbxl5*^F/F^ (control) and *Alb-*Cre*/Fbxl5*^F/F^ (FBXL5 liver-KO) mice at 3 h after iron administration. FDR-q, false discovery rate q value; NES, normalized enrichment score.

### Integrated ferroptosis response gene signature dictates ferroptosis-related pathological response in mouse liver

Ferroptosis, induced by an imbalance between iron toxicity and antioxidants, can also be influenced by a reduction in redox potential [18]. Among such mouse models, APAP overdose-induced GSH depletion serves as a widely used mouse model for ferroptosis induction in the liver [19]. To better understand the hepatic gene signature indicative of ferroptosis induction *in vivo*, we aimed to narrow down the gene list obtained from our iron overload-induced ferroptosis model by comparing it with that from the APAP-induced ferroptosis model. First, we characterized the genes whose expressions are commonly upregulated in APAP overdose-induced and iron overload-induced ferroptosis models (Fig. 3A). Re-analysis of gene expression data from Zhang et al. [29] identified 479 DEGs (220 and 259 genes with upregulated and downregulated expressions, respectively) (Table S3) in the livers of APAP-treated mice compared to that in control mice (Fig. 3B, left). Furthermore, RNA sequencing analysis identified 1,242 DEGs (733 and 509 genes with upregulated and downregulated expressions, respectively) (Table S4) in the livers of FBXL5 liver-KO mice at 0 and 3 h after iron administration (Fig. 3B, right). The top 100 genes with upregulated expressions common to both models were selected to establish a gene set termed the “integrated gene signature for hepatic ferroptosis response (iFerroptosis)” (Fig. 3C and Table S5). Next, we investigated whether the iFerroptosis gene set could better characterize the pathological features of ferroptosis in the mouse liver than the existing gene set for ferroptosis (mmu04216). We applied the iFerroptosis gene list to the liver transcriptome of IRI (Table S6) [29], metabolic dysfunction-associated steatohepatitis (MASH) (Table S7) [30], and CCl_4_-induced acute liver injury (Table S8) [31]. GSEA using the iFerroptosis gene list revealed significant enrichment of hepatic ferroptosis response genes in the IRI (Fig. 3D) and MASH (Fig. 3E) models (*p* <□0.001). This is consistent with previous findings showing that ferroptosis is associated with the pathogenesis of IRI and MASH in mice [17]. In contrast, the application of the existing gene set (mmu04216) did not achieve statistical significance in the enrichment of ferroptosis-related genes in the liver transcriptomes of IRI and MASH (*p*□=□0.92 and *p*□=□0.49, respectively). Given that mmu04216 gene set is composed of genes associated with the cellular pathway of ferroptosis induction, these results suggest that the iFerroptosis gene list, which dictates the tissue-level response to ferroptosis, may better identify the pathogenic features of ferroptosis under multiple liver pathogenesis scenarios. Notably, GSEA using the iFerroptosis gene list did not show statistically significant enrichment of hepatic ferroptosis response genes in the liver transcriptome of CCl_4_-induced acute liver injury (*p* = 0.97) (Fig. 3F), suggesting that the iFerroptosis gene list predominantly detects ferroptosis-associated pathogenesis in the murine liver.

**Fig. 3.**
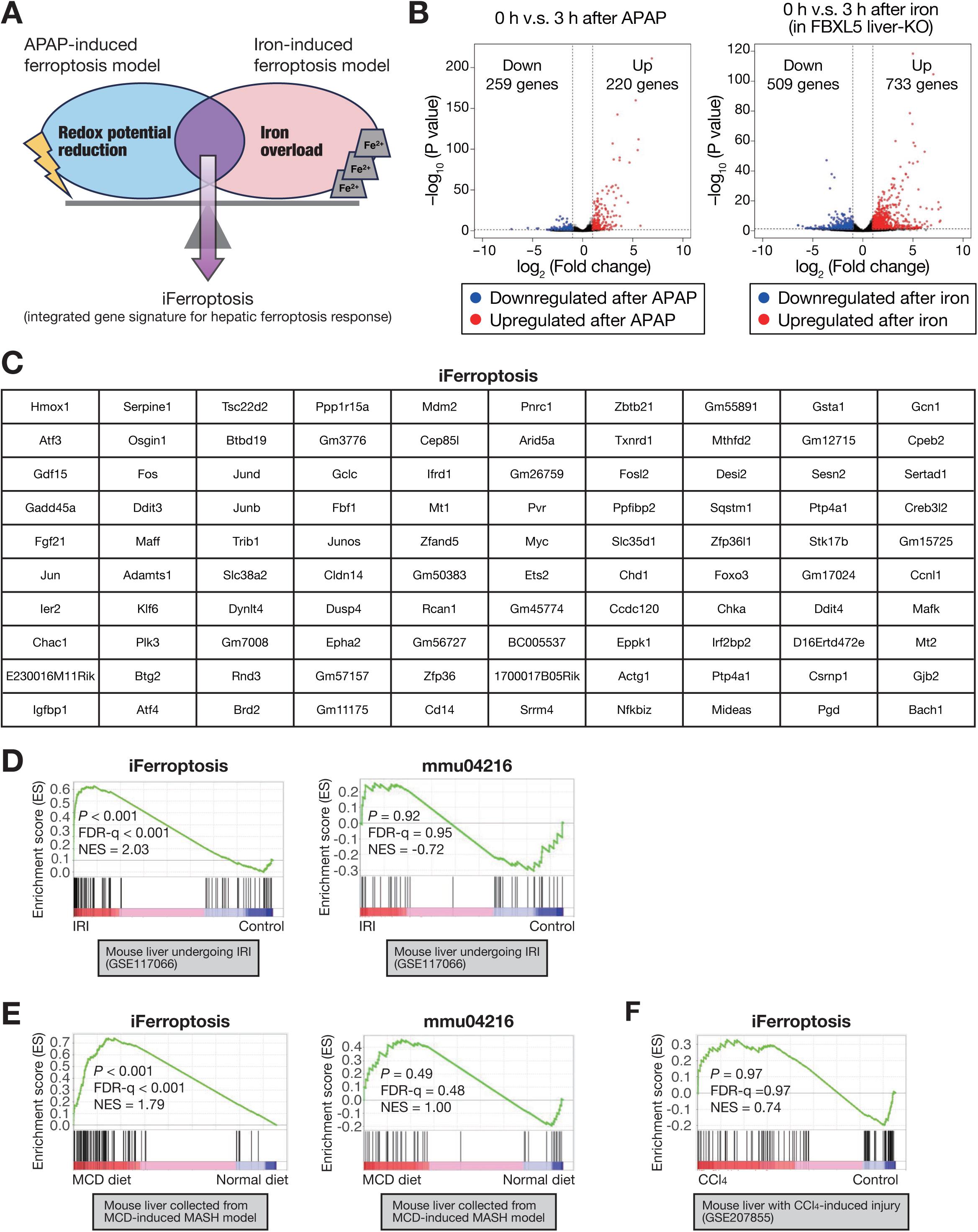
Integrated ferroptosis response gene signature dictates ferroptosis-related pathological response in mouse liver. (A) Schematic overview of the strategy for creating integrated gene signature for hepatic ferroptosis response (iFerroptosis). (B) Left panel shows volcano plot of differentially expressed genes (DEGs; fold change > 2 and adjusted *p* < 0.01) in the liver of mice treated with acetaminophen (APAP), comparing samples collected at 0 h and 3 h after intrapreneurial injection. Right panel shows DEGs in the liver of *Alb-*Cre*/Fbxl5*^F/F^ (FBXL5 liver-KO) mice treated with iron, comparing samples collected at 0 h and 3 h after oral gavage injection. The genes with upregulated and downregulated expressions are shown in red and blue, respectively. RNA sequencing data for the APAP-induced ferroptosis model was obtained from Li et al. [42]. (C) The top 100 genes with upregulated expressions in the iFerroptosis gene signature. (D) GSEA of the iFerroptosis gene signature (left panel) or the KEGG ferroptosis pathway (mmu04216) (right panel) in the mouse liver undergoing ischemia-reperfusion injury (IRI). RNA sequencing data were obtained from Zhang et al. [29]. FDR-q, false discovery rate q value; NES, normalized enrichment score. (E) GSEA of the iFerroptosis gene signature (left panel) or the KEGG ferroptosis pathway (mmu04216) (right panel) in the liver of mouse fed with methionine–choline-deficient (MCD) diet to mimic metabolic dysfunction-associated steatohepatitis (MASH) progression. RNA sequencing data was obtained from Li et al. [30]. (F) GSEA of the iFerroptosis gene signature in the mouse liver with carbon tetrachloride-induced injury. RNA sequencing data was obtained from Xi et al. [31].

### Iron overload aggravates hepatic IRI in mice

Given that GSEA using the iFerroptosis gene list suggested an association between ferroptosis and the pathogenesis of IRI in the murine liver, we investigated whether disruption of iron homeostasis promotes ferroptosis and exacerbates IRI. To this end, control and FBXL5 liver-KO mice were fasted for 12–16 h, followed by 70% hepatic ischemia for 30 min and reperfusion for 3 h (Fig. 4A). Serum levels of AST, ALT, and LDH increased in FBXL5 liver-KO mice after IRI compared to those in controls, and these effects were significantly suppressed by Fer-1 treatment (Fig. 4B). Consistent with these results, microscopic analysis of sections stained with 4-HNE suggested an increase in lipid peroxidation in the livers of FBXL5 liver-KO mice, which was mitigated by Fer-1 treatment (Fig. 4C). These results demonstrated that iron overload in FBXL5 liver-KO mice promoted ferroptosis during IRI, leading to severe liver damage. We next validated the iFerroptosis gene signature in this context by individually evaluating the expression of iFerroptosis genes, including heme oxygenase 1 (*Hmox1*), growth differentiation factor 15 (*Gdf15*), fibroblast growth factor 21 (*Fgf21*), and immediate early response2 (*Ier2*) (Fig.4D). RT and quantitative PCR analyses revealed that the expression of these genes was substantially upregulated under IRI in the livers of *Fbxl5* liver-KO mice, which was reduced by Fer-1 treatment (Fig. 4D). These results suggest that the expression levels of iFerroptosis genes are associated with the induction of ferroptosis during IRI in mice, prompting us to apply this list to the human IRI transcriptome. Indeed, GSEA using the iFerroptosis gene list revealed that human livers undergoing IRI during conventional liver transplantation (Table S9) [32] displayed significant enrichment of hepatic ferroptosis response genes (*p*□<□0.001) (Fig. 4E), suggesting that ferroptosis is associated with the pathogenesis of hepatic IRI in humans. Taken together, our data indicate the potential of the iFerroptosis gene list to predict ferroptosis-associated pathogenesis in both human and mouse livers.

**Fig. 4.**
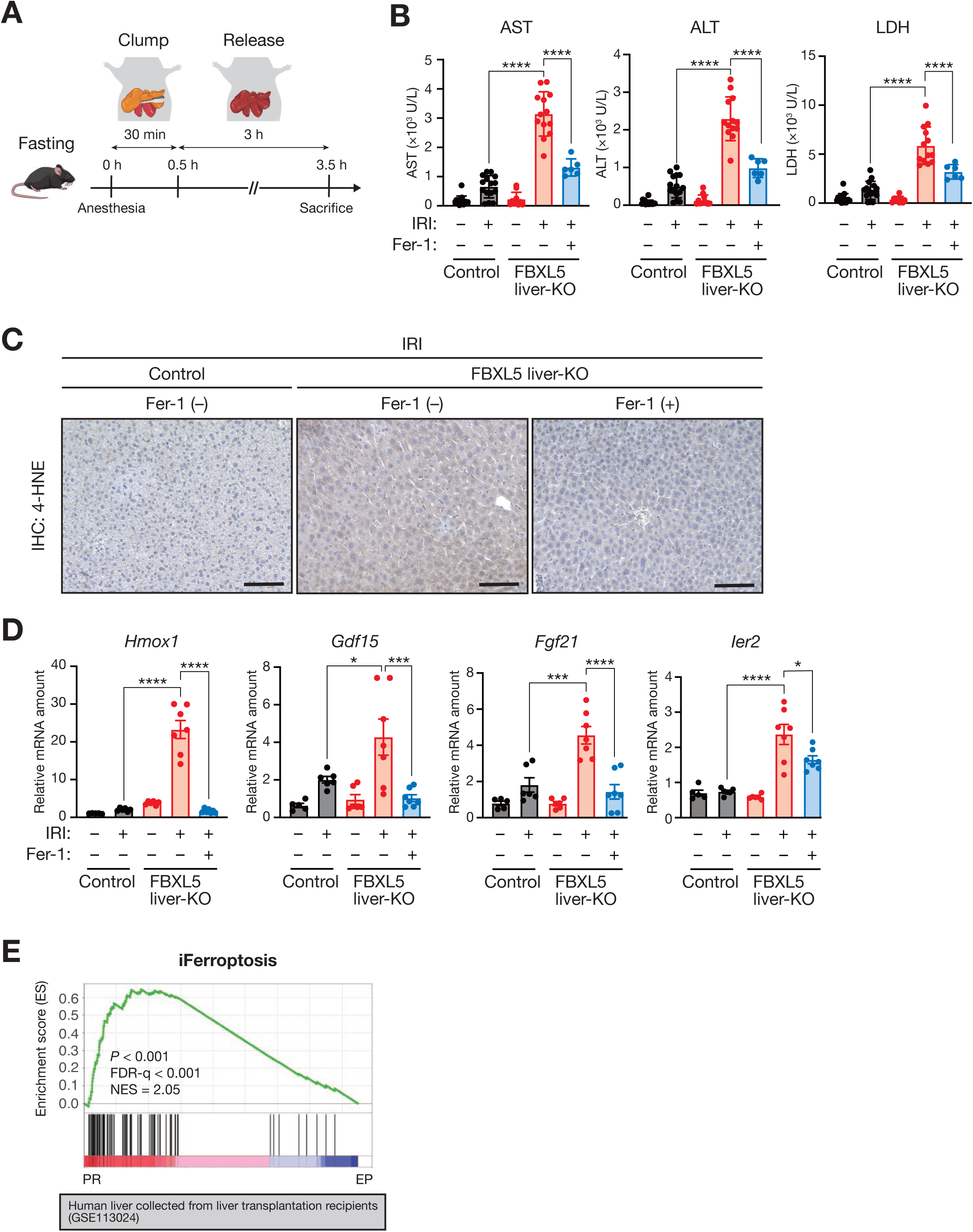
Iron overload aggravates hepatic ischemia-reperfusion injury in mice. (A) Schematic overview of the experimental procedure for 70% IRI model. (B) Serum levels of AST, ALT and LDH in *Fbxl5*^F/F^ (control) and *Alb-*Cre*/Fbxl5*^F/F^ (FBXL5 liver-KO) mice after IRI. Where indicated, FBXL5 liver-KO mice were provided (or not) with Fer-1 for 1 h prior to IRI. Data are represented as mean ± SD. The number of mice used for this data are as follows: n = 16 and 15 mice for sham and IRI, respectively, for the control group; and n = 10, 13, 6 mice for sham, IRI, IRI with Fer-1 treatment, respectively, for the FBXL5 liver-KO group. ∗∗∗∗*p* < 0.0001 (one-way ANOVA test followed by Tukey’s multiple comparison test). (C) IHC staining with antibodies against 4-HNE of the liver of *Fbxl5*^F/F^ (control) and *Alb-*Cre*/Fbxl5*^F/F^ (FBXL5 liver-KO) mice after IRI. Mice were provided (or not) with Fer-1 as described in (B). Scale bars represent 100 μm. (D) RT and real-time PCR analysis of the indicated mRNA in liver tissues of *Fbxl5*^F/F^ (control) and *Alb-*Cre*/Fbxl5*^F/F^ (FBXL5 liver-KO) mice after IRI. Mice were provided (or not) with Fer-1 as described in (B). The expression levels of the following genes were studied: heme oxygenase 1 (*Hmox1*); growth and differentiation factor 15 (*Gdf15*); fibroblast growth factor 21 (*Fgf21*); immediate early response 2 (*Ier2*). Data are represented as mean ± standard error of mean (SEM). The number of mice used for this data are as follows: n = 5 and 6 mice for sham and IRI, respectively, for the control group; and n = 6, 7, 7 mice for sham, IRI, IRI with Fer-1 treatment, respectively, for the FBXL5 liver-KO group. ∗*p* < 0.05; ∗∗∗*p* < 0.001; ∗∗∗∗*p* < 0.0001 (one-way ANOVA test followed by Tukey’s multiple comparison test). (E) GSEA of the iFerroptosis gene signature in the transplanted human liver, comparing samples collected at the end of preservation and 1 h post-graft revascularization during conventional liver transplantation. RNA sequencing data was obtained from Guo et al. [32].

### Patients with high iron status show sustained post-operative liver damage

Our data suggest that iron aggravates hepatic IRI. IRI is often a problem not only in liver transplantation but also in liver resection surgery [34, 35]. The Pringle maneuver method, used to prevent bleeding during liver resection, causes temporary ischemia and subsequent reperfusion of the liver [36, 37]. Therefore, we investigated the impact of iron and ferroptosis on recovery from IRI caused by hepatectomy in humans. Pre-operative serum iron levels were measured in 174 of 1,014 liver resections performed for hepatocellular carcinoma at Kumamoto University between 2007 and 2023, where the Pringle maneuver method was used to prevent intraoperative hemorrhage. These 174 cases were divided into two groups based on pre-operative serum iron levels (cutoff value: 150 µg/dL): a low-iron group (n = 143) and a high-iron group (n = 31) (Fig. 5A and Table 1). This criterion allowed these groups to be further divided according to serum iron levels, with AST and ALT levels increasing in the high-serum iron group (Fig. 5B). These results suggest that elevated iron levels in patients with hepatocellular carcinoma are associated with increased liver damage. In contrast, there was no significant difference in albumin levels and indocyanine green retention 15 (ICG R15) values between the low- and high-iron groups, which were used to evaluate resectability for hepatectomy (Table 1). Additionally, there was no significant difference in the duration of the Pringle maneuver between the two groups, suggesting that the amount of ischemia-reperfusion stress was comparable in both groups (Fig. 5B).

**Fig. 5.**
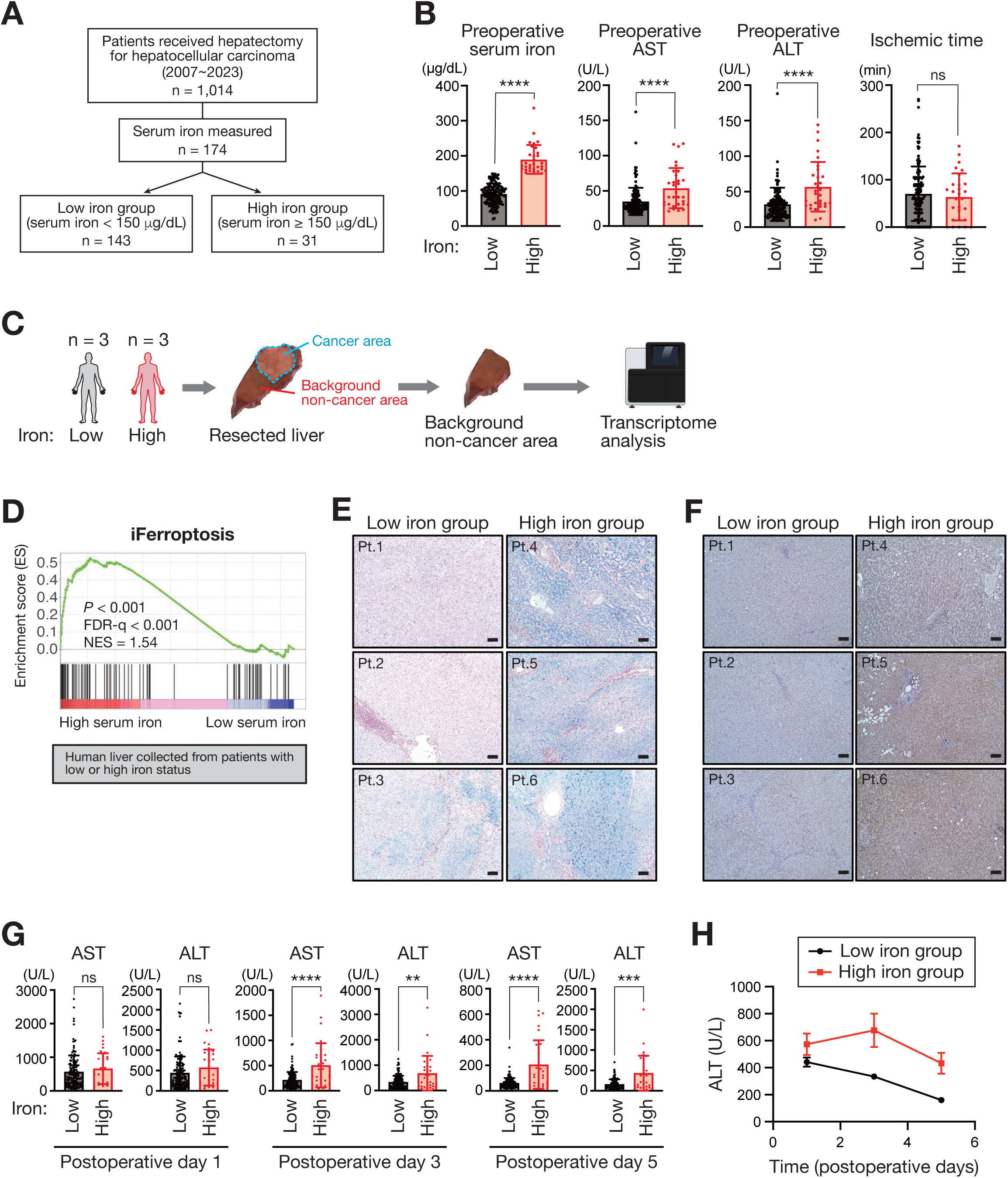
Patients with high iron status show sustained post-operative liver damage. (A) Schematic flow chart of clinical sample collection. (B) Pre-operative serum levels of iron, AST, and ALT, as well as ischemic time during operation, in patients with hepatocellular carcinoma with low- or high-iron status. Data are represented as mean ± SD. The number of patients are as follows: serum levels of iron, AST, and ALT, n = 143 and 31 patients for low- and high-iron group, respectively; ischemic time, n = 138 and 28 patients for low- and high-iron group, respectively. ns, not significant (*p* > 0.05); ∗∗∗∗p < 0.0001 (Mann-Whitney U-test). (C) Schematic overview of patient sample preparation for the transcriptome analysis. (D) GSEA of the iFerroptosis gene signature in the resected human liver, comparing samples collected from patients with low- and high-iron status. (E and F) Histological analysis of resected non-cancerous human liver. Perls Prussian blue staining for the detection of iron is shown in (E), IHC staining with antibodies against 4-HNE is shown in (F). Scale bars represent 100 μm. (G) Serum levels of AST and ALT in patients with hepatocellular carcinoma with low- or high-iron status at the indicated times after hepatectomy. Data are represented as mean ± SD; n = 143 and 31 patients for low- and high-iron group, respectively. ns, not significant (*p* > 0.05); ∗∗*p* < 0.01; ∗∗∗*p* < 0.001; ∗∗∗∗*p* < 0.0001 (Mann-Whitney U-test). (H) The time course of changes in serum ALT levels in patients, as described in (G), is represented as mean ± SEM data.

**Table 1.** Association between serum iron levels and clinicopathological factors in patients with hepatocellular carcinoma.

|  | Low iron < 150 µg/dL (n=143) | High iron ≥ 150 µg/dL (n=31) | *p*-value |
| --- | --- | --- | --- |
| Age | 69.7±9.4 | 66.3±11.2 | 0.10 |
| Sex (M / F) | 102/41 | 24/7 | 0.41 |
| BMI (kg/m<sup>2</sup>) | 23.5± | 24.2±2.7 | 0.31 |
| HBs-Ag (+) | 31(22%) | 4(13%) | 0.38 |
| HCV-Ab (+) | 59(41%) | 20(65%) | 0.03 |
| Albumin (g/dL) | 4.1±0.4 | 4.0±0.4 | 0.74 |
| AST (IU/L) | 36.9±21 | 56.0±28 | <0.001 |
| ALT (IU/L) | 34.5±25 | 61.3±35 | <0.001 |
| CRP (mg/dL) | 0.14±0.40 | 0.084±0.08 | 0.46 |
| ICG R 15min (%) | 13.0±7.1 | 13.3±7.6 | 0.81 |
| Prothrombin activity (%) | 96±19 | 97±24 | 0.82 |
| Plt (10<sup>4</sup>/µL) | 18.2±18.3 | 14.3±3.9 | 0.26 |
| Tumor size (mm) | 27.5±11.8 | 24.3±8.3 | 0.18 |
| Tumor number | 1(1-3) | 1(1-3) | 0.35 |
| AFP (mAU/mL) | 905±3804 | 126±373 | 0.28 |
| AFP-L3 (%) | 12±21.5 | 7.7±12.4 | 0.32 |
| PIVKA-II (mAU/mL) | 189±450 | 335±831 | 0.21 |
| Operation time (min) | 373±110 | 398±97 | 0.29 |
| Blood loss (mL) | 457±440 | 378±323 | 0.37 |
| Ischemic time (min) | 71±58 | 64±50 | 0.56 |
| Laparoscopic resection | 50(35%) | 12(39%) | 0.67 |
| Anatomic resection | 81(57%) | 18(58%) | 0.97 |

Abbreviation: BMI, body mass index; HBV, hepatitis B virus; HCV, hepatitis C virus; AST, aspartate aminotransferase; ALT, alanine aminotransferase; CRP, C reactive protein; ICG R 15min, indocyanine retention rate at 15 min; PT, prothrombin time; Plt, platelet count; AFP, $\alpha$ -fetoprotein, AFP-L3, Lens culinaris agglutinin-A-reactive $\alpha$ -fetoprotein, PIVKA-II, protein induced by vitamin K absence or antagonist-II

To investigate the mechanisms underlying the higher pre-operative transaminase levels in the high-iron group than in the low-iron group, we collected three liver samples from each group and examined the gene expression profiles of non-cancerous background regions of the liver (Fig. 5C and Table S10). GSEA using the iFerroptosis gene list revealed a significant enrichment of the hepatic ferroptosis response in patients with a high iron status compared to those with a low iron status (Fig. 5D). Histological analysis revealed extensive accumulation of iron (Fig. 5E) and lipid peroxidation (Fig. 5F) in the high-iron group. These observations suggest that patients with high serum iron levels accumulate iron in the liver, thereby increasing the susceptibility of hepatocytes to ferroptosis and resulting in liver damage. Next, we investigated the impact of iron accumulation on recovery from hepatectomy-induced hepatic IRI. The serum levels of AST and ALT were high and showed no statistical difference between the low- and high-iron groups one day after surgery, possibly reflecting massive liver destruction because of hepatectomy (Fig. 5G). This increase in serum transaminase levels gradually decreased over time in the low-iron group, as evident from the AST and ALT levels on post-operative days 3 and 5 (Figs. 5G and 5H). In contrast, patients with high iron status showed delayed recovery from hepatectomy, with a significant difference in the serum levels of AST and ALT between the low-iron and high-iron groups (Figs. 5G and 5H). Therefore, our data indicate that patients with high iron levels accumulate iron in the liver, disrupting the prompt recovery from hepatic IRI by increasing the susceptibility of hepatocytes to ferroptosis.

## DISCUSSION

Identifying and characterizing ferroptosis-related pathogenesis in liver disease using bulk RNA sequencing data has posed challenges for several reasons. These include context-dependent involvement of ferroptosis-related molecules and the lack of a consistent gene set capable of reliably identifying a ferroptosis signature *in vivo*. In the present study, we developed a hepatic ferroptosis response gene set (iFerroptosis) comprising 100 genes whose expressions are commonly upregulated in iron- and APAP-induced ferroptosis models. Validation was conducted in murine models of IRI and MASH, as well as in human liver samples obtained during resections for hepatocellular carcinoma. To construct this gene set, we established a novel mouse model of iron overload-induced ferroptosis using liver-specific *Fbxl5* knockout mice. Using publicly available RNA sequencing data, we demonstrated the applicability of the iFerroptosis gene set in both murine and human context, emphasizing the pivotal role of iron and ferroptosis in ischemic liver injury. The proposed iFerroptosis gene signature encompasses transcriptional profiles not only from hepatocytes but also from other cells involved in tissue responses, including cholangiocytes, stellate cells, endothelial cells, Kupffer cells, and immune cells. This contrasts with the existing ferroptosis-related gene set (mmu04216), which comprises genes associated with the cellular pathways involved in ferroptosis induction. Notably, our iFerroptosis gene signature does not include genes known for their critical roles in ferroptosis induction, such as *GPX4* and acyl-CoA synthetase long-chain family member 4 (*ASCL4*) [4]. Pathway gene sets, such as mmu04216, typically compile genes linked to the pathway from individual studies or large-scale analyses. Although these pathway gene lists may capture events that could be associated with ferroptosis under certain conditions, the complex response of the liver to ferroptosis *in vivo* suggests that our newly constructed gene list may more accurately depict ferroptosis features in multiple liver pathogenesis scenarios. However, this study exclusively focused on hepatic responses to ferroptosis; thus, it is uncertain whether the iFerroptosis gene set is applicable across other tissues. Future studies could extend this experimental approach by generating tissue-specific *Fbxl5* knockout mice in other organs, such as the kidneys and heart, and developing gene signatures that describe tissue-specific responses to iron-induced ferroptosis.

Liver resection is the optimal curative treatment for hepatobiliary malignancies, such as liver cancer. For decades, the Child-Pugh score has been utilized to assess liver function. This assessment incorporates specific blood markers, such as serum bilirubin, albumin, and prothrombin levels, along with clinical observations, including the presence of hepatic encephalopathy and ascites [38]. In recent years, the utility of the ALBI (albumin-bilirubin) score, which uses albumin and bilirubin levels, has been reported. Additionally, the indocyanine green retention rate at 15 min (ICG-R15), which measures indocyanine green, has been reported to correlate with liver reserve function as an assessment parameter [39]. In the present study, we demonstrated that pre-operative serum iron levels are associated with perioperative liver damage in patients with hepatocellular carcinoma. Therefore, incorporating the pre-operative serum iron status into these scoring criteria could be beneficial for preventing post-operative complications and liver failure. Our findings demonstrate that iron overload induced by the liver-specific deletion of *Fbxl5* aggravates hepatic IRI in mice, suggesting that strict control of hepatic iron status is crucial for managing hepatic IRI. As IRI is a notable clinical problem, especially following liver transplantation and major resection [40, 41], targeting iron in the liver may be a promising strategy for mitigating IRI and improving liver surgery outcomes.

In conclusion, our study identified a gene signature indicative of hepatic ferroptosis in both mice and humans. Utilizing the iFerroptosis gene set may help to further understand ferroptosis-related hepatic disorders and offer insights into potential therapeutic strategies targeting ferroptosis to improve outcomes in patients with liver diseases.

## Abbreviations

FBXL5: F-box and leucine-rich repeat protein 5
IRI: ischemia-reperfusion injury
GSH: glutathione
GPX4: glutathione peroxidase 4
APAP: acetaminophen
PUFA: polyunsaturated fatty acid
PRM: phosphatidylcholine
IRP1: iron regulatory protein 1
PBS: phosphate-buffered saline
RT: reverse transcription
GSEA: gene set enrichment analysis
AST: aspartate aminotransferase
ALT: alanine aminotransferase
LDH: lactate dehydrogenase
4-HNE: 4-hydroxynonenal
DMSO: dimethyl sulfoxide
PCA: principal component analysis
DEGs: differentially expressed genes
MASH: metabolic dysfunction-associated steatohepatitis
CCL4: carbon tetrachloride4
HMOX1: heme oxygenase 1
GDF15: growth differentiation factor 15
FGF21: fibroblast growth factor 21
IER2: immediate early response2
qPCR: quantitative polymerase chain reaction
ACSL4: acyl-CoA synthetase long chain family member 4
ICGR15: indocyanine green retention 15
ALBI: albumin-bilirubin
PC: phosphatidylcholine
MS/MS: tandem mass spectrometry
HRMS: high-resolution mass spectrometry
LC/HRMS: liquid chromatography coupled with HRMS
MALDI-MS/MS/MSI: matrix-assisted laser desorption/ionization-tandem MS-MS imaging
ANOVA: analysis of variance
DDBJ: DNA Data Bank of Japan
FAC: ferric ammonium citrate
H&E: hematoxylin and eosin
KEGG: Kyoto Encyclopedia of Genes and Genomes
NES: normalized enrichment score
SD: standard deviation
SEM: standard error of mean

## Data availability statement

The scRNA-seq data generated in this study have been deposited in the DNA Data Bank of Japan (DDBJ) at PRJDB18097 and PRJDB18207. All other raw data generated in this study are available from the corresponding author upon request.

## Acknowledgements

This study was supported by Japan Society for the Promotion of Science (JSPS) KAKENHI grants (23K18098, 24H00864, and 24H00865 to T.Moroishi), Japan Science and Technology Agency (JST) FOREST Program (JPMJFR226J to T.Moroishi), Kobayashi Foundation for Cancer Research (to T.Moroishi), Princess Takamatsu Cancer Research Fund (to T.Moroishi), Ichiro Kanehara Foundation (to T.Moroishi), Foundation for Promotion of Cancer Research in Japan (to T.Moroishi), Kato Memorial Bioscience Foundation (to T.Moroishi), Japan Agency for Medical Research and Development (AMED; JP23fk0210103 to Y.T.), and Collaborative Research Grant from Center for Metabolic Regulation of Healthy Aging (CMHA; to Y.T.). The authors thank Core Laboratory for Medical Research and Education, Kumamoto University School of Medicine, for their technical support.

---

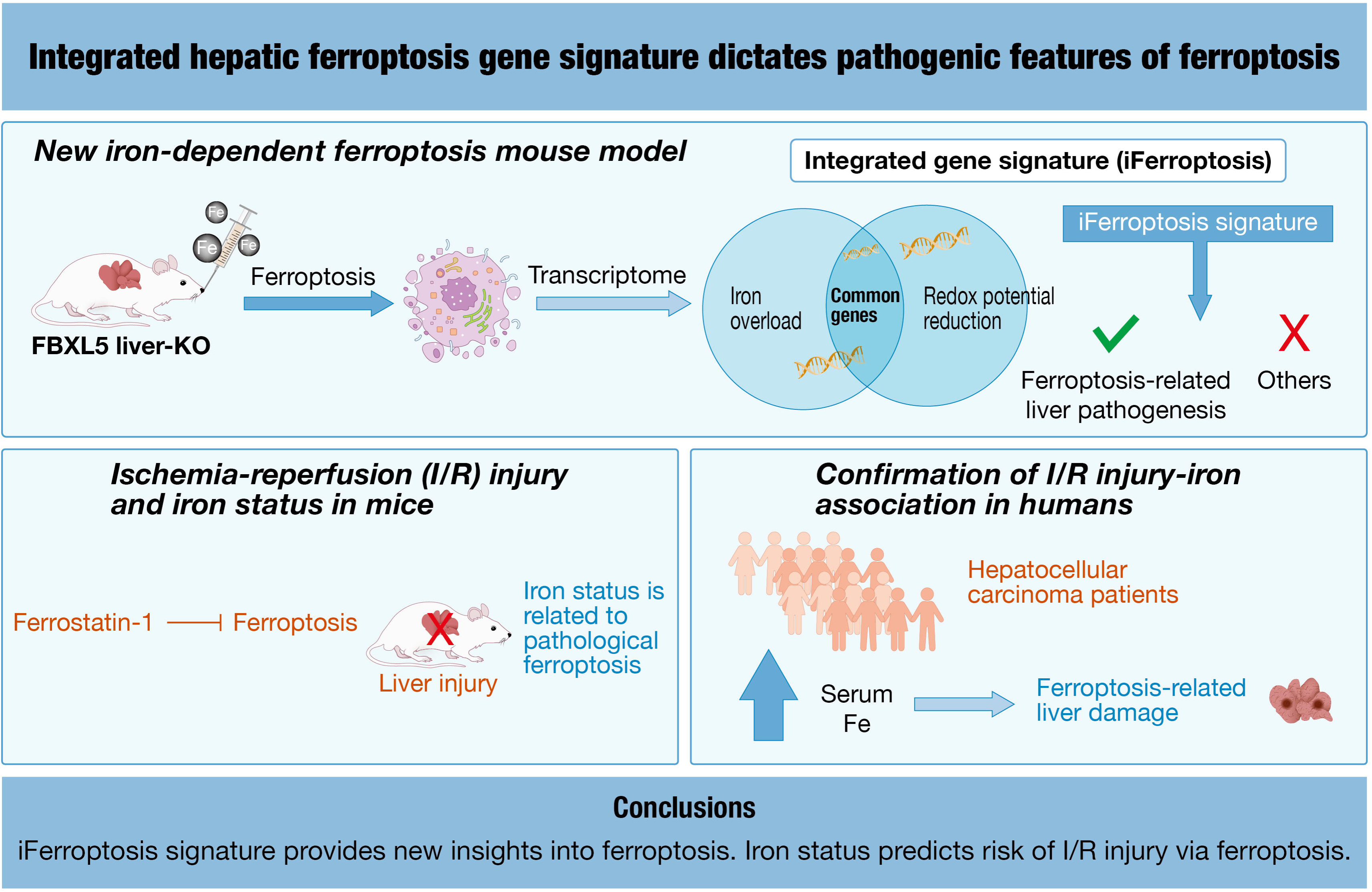

